# Photometallobiocatalytic Asymmetric Radical-Mediated Cross-Coupling of Organotrifluoroborate Salts and Pyridotriazoles

**DOI:** 10.64898/2026.08.11.744224

**Authors:** Huanan Wang, Binh Khanh Mai, Xingjie Zhang, Chongtao Li, Peng Liu, Yang Yang

## Abstract

The cooperative integration of photoredox catalysis and metalloenzyme catalysis has emerged as a powerful strategy for enabling stereoselective radical transformations beyond the capabilities of either catalytic mode alone. Herein, we report a photometallobiocatalytic enantioselective intermolecular C–C cross-coupling of pyridotriazoles and secondary alkyltrifluoroborate salts through cooperative catalysis between an organic photosensitizer and an engineered protoglobin. By combining visible-light-mediated radical generation with enzymatic activation of pyridotriazoles to form reactive Fe carbenoid intermediates, this transformation enabled highly enantioselective radical C–C bond formation through a proposed outer-sphere coupling mechanism. Through biocatalyst mining and directed evolution, engineered Aeropyrum pernix protoglobin catalysts were developed that catalyzed this radical C–C coupling with excellent efficiency and stereocontrol. The photobiocatalytic platform exhibited a broad substrate scope with respect to both secondary alkyltrifluoroborate salts and pyridotriazoles, affording a range of valuable N-heterocyclic products in excellent yields and enantioselectivities. Mechanistic studies supported the involvement of radical intermediates and revealed spontaneous binding between the photocatalyst eosin B and the engineered metalloenzyme. By leveraging cooperative photometallobiocatalysis, this work established an underexplored strategy for asymmetric intermolecular radical cross-coupling via an outer-sphere mechanism, further expanding the catalytic repertoire of transition-metal carbenoid chemistry.

**Entry for the Table of Contents:** An enantioselective photometallobiocatalytic cross-coupling of pyridotriazoles and secondary alkyltrifluoroborate salts is developed. Cooperative catalysis using eosin B and an engineered protoglobin combines visible-light-mediated radical generation with enzymatic metal carbenoid activation, affording valuable *N*-heterocyclic products in excellent yield and enantioselectivity through an outer-sphere radical coupling pathway.

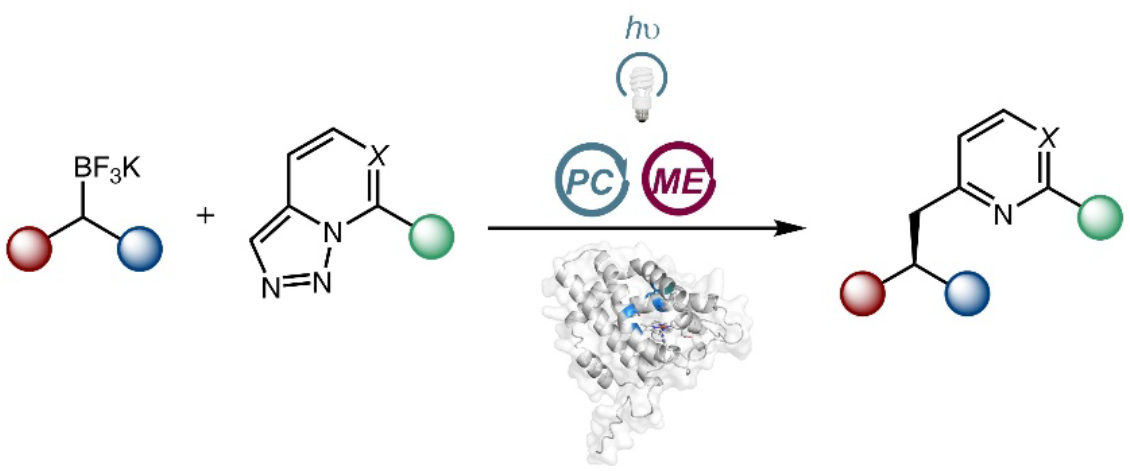

## Introduction

The cooperative merger of photoredox catalysis and metalloenzyme catalysis has recently emerged as a powerful strategy for developing asymmetric radical transformations that are difficult to achieve using either catalytic mode alone. In this new mode of cooperative photometallobiocatalysis, the visible-light photocatalyst enables mild and controlled generation of transient free radical intermediates from readily available organic precursors,^1,2^ while the metalloenzyme facilitates the efficient formation of highly reactive organometallic intermediates within the protein’s confined active site, ultimately permitting challenging C–C and C–heteroatom cross-couplings to occur with excellent stereoselectivity.^3,4^ In particular, recent studies from our laboratory and others have demonstrated the efficient coupling of radical species generated outside the protein scaffold, thereby addressing the long-standing challenge of achieving catalytic stereocontrol in intermolecular free radical reactions. ^5–10^ Collectively, these desirable features establish cooperative photometallobiocatalysis as a promising platform for expanding the synthetic utility of metalloenzymes beyond their native functions, ushering in a range of new-to-nature stereoselective reactions.^10^

Inspired by cooperative photobiocatalytic reactions using cofactor-dependent and independent enzymes developed by our group and other researchers, ^5–10^ we initiated a program leveraging cooperative photobiocatalysis to unlock novel radical C–C cross-coupling reactivity of transition-metal carbenoids. Over the past several decades, groundbreaking studies in metal carbenoid chemistry based on Rh, Co, Fe and other transition metals have led to a plethora of stereoselective C–C bond forming reactions.^11^ Furthermore, through the engineering of naturally occurring hemoproteins, pioneering studies by Arnold, Fasan, Hartwig and others have culminated in a diverse array of biocatalytic C–C bond forming reactions via a carbene transfer mechanism.^12,13^ Despite this progress, the productive merger of transition-metal carbenoid chemistry with photoredox radical catalysis has remained elusive until our recent report.

**Scheme 1.**
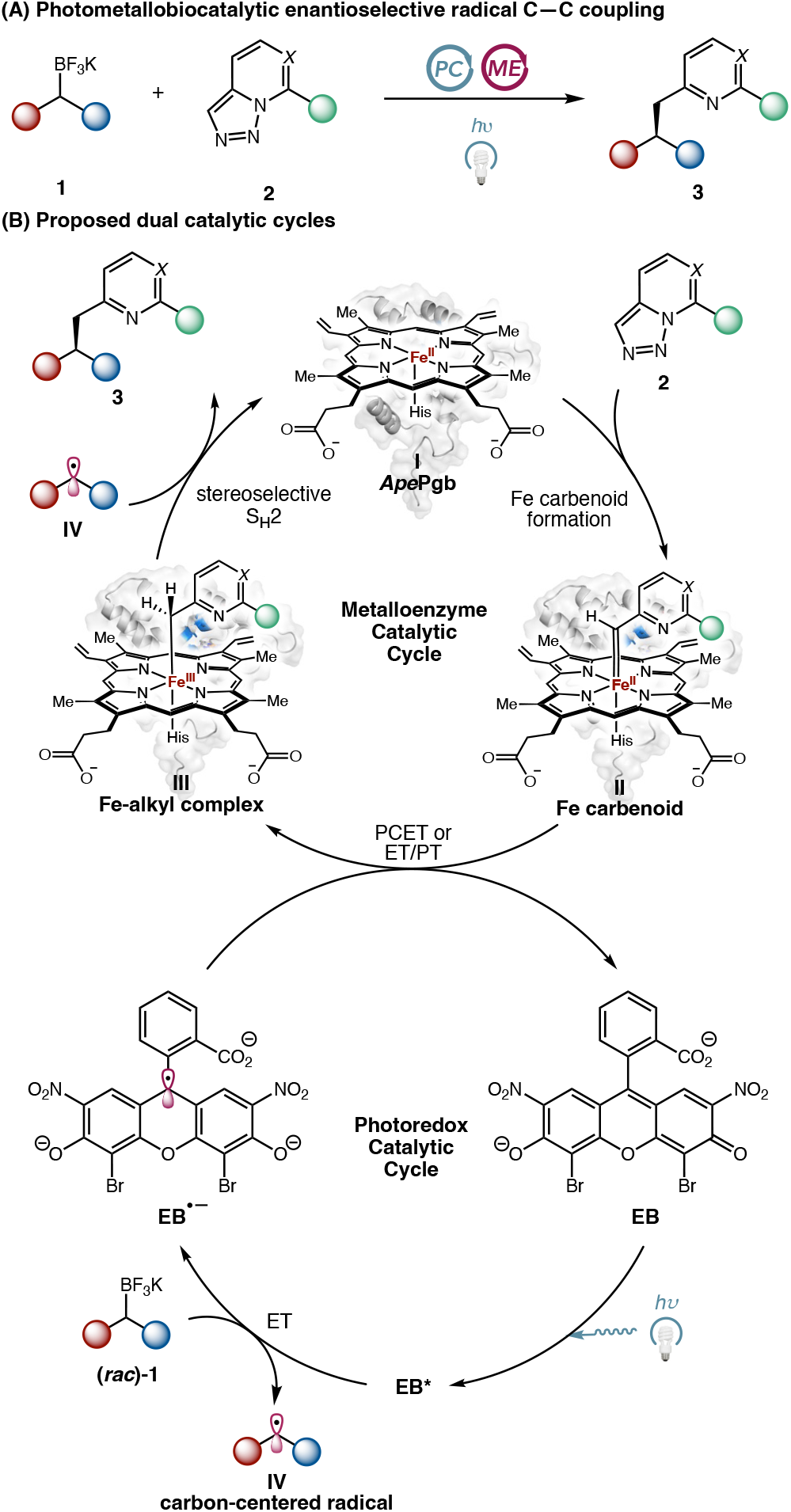
Mechanistic study for photometallobiocatalytic asymmetric cross-coupling of benzyltrifluoroborate salts and pyridotriazoles. Heme was prepared as a 0.5 M NaOH stock solution for use in studies detailed herein.

In 2025, enabled by cooperative photometallobiocatalysis, our laboratory introduced a new class of formal metal carbenoid– radical cross-coupling reactions.^14^ By capitalizing on the diradical reactivity of Fe carbenoids, this photobiocatalytic strategy enabled outer-sphere radical C–C coupling to occur at the carbenoid carbon with excellent efficiency. Furthermore, through directed evolution of cytochromes *c*, we achieved asymmetric cross-couplings of α-diazocarbonyl compounds and benzyltrifluoroborate salts. Despite the excellent synthetic potential of this new radical coupling mechanism, our prior study largely focused on the asymmetric construction of stereogenic centers at the α-position of Fe-carbenoid intermediates through an enzyme-controlled enantioselective protonation mechanism.^14^ In contrast, the enantioconvergent cross-coupling of secondary alkyl radical precursors, particularly conformationally flexible acyclic substrates, remains underexplored.

Based on these recent findings, we questioned whether this intermolecular photobiocatalytic metal carbenoid–radical cross-coupling could be generalized for the stereocontrolled synthesis of valuable nitrogen heterocyclic compounds. In this context, we became interested in photobiocatalytic cross-coupling of pyridotriazoles, a recently emerged class of readily accessible carbene precursors,^15,16^ for the asymmetric construction of stereogenic centers bearing heterocyclic substituents, which represent ubiquitous structural motifs in bioactive compounds and pharmaceuticals.^17^ The proposed cross-coupling of pyridotriazoles and alkylboron reagents, including its racemic version, was not previously known in either organic chemistry or enzymology. Thus, the successful implementation of this photometallobiocatalytic transformation would further expand the boundaries of both organic synthesis and biocatalysis. Important prior studies from the Fasan group demonstrated the utility of pyridotriazoles in myoglobin-catalyzed stereoselective cyclopropanation and C–H insertion reactions.^18^ Nevertheless, further application of these N-heterocyclic substrates in a wider range of asymmetric C–C bond forming reactions remains underexplored. We were particularly interested in the enantioconvergent photobiocatalytic coupling of racemic secondary alkylboron reagents with pyridotriazoles, as these transformations would require an enantioinduction mechanism distinct from the asymmetric protonation pathway reported previously by our group. We postulated that mining structurally diverse naturally occurring hemoproteins, particularly thermostable protoglobins, could identify an excellent protein scaffold for subsequent biocatalyst optimization. Furthermore, we envisioned that directed evolution combined with high-throughput photobiocatalysis would furnish heme-dependent biocatalysts with substantially improved stereoselectivity, thereby affording a general strategy for asymmetric intermolecular cross-couplings to access valuable N-heterocyclic products.

Herein, we report a photometallobiocatalytic enantioselective coupling of benzyltrifluoroborate salts and pyridotriazoles through cooperative substrate activation using an organic photosensitizer and an evolved *Aeropyrum pernix* protoglobin (*Ape*Pgb, Scheme 1(A)). A plausible mechanism for the cooperative photometallobiocatalytic C–C coupling is illustrated in Scheme 1(B). In the proposed photoredox cycle, visible-light excitation of eosin B (EB) would generate the excited-state EB*, which serves as a potent photooxidant. Single-electron oxidation of the organotrifluoroborate substrate (**1**) with EB* would produce a transient prochiral carbon-centered radical and the reduced photocatalyst EB^•−^. Concurrent to the photoredox cycle, within the metalloenzyme cycle, an appropriately engineered *Ape*Pgb variant (**I**) would activate the pyridotriazole substrate (**2**) to form an iron carbenoid intermediate (**II**). Subsequent electron transfer (ET) from EB^•−^ to the enzymatic iron carbenoid, coupled with proton transfer (PT), would convert **II** into an Fe–alkyl species (**III**). Finally, the prochiral secondary alkyl radical would undergo an enantioselective bimolecular homolytic substitution (S_H_2) reaction^19^ with the Fe–alkyl intermediate **III**, furnishing C–C coupling product **3** in an enantioenriched fashion.

## Results and Discussion

At the commencement of our studies, using benzyltrifluoroborate salt **1a** and pyridotriazole **2a** as model substrates and eosin Y as the photosensitizer, we evaluated an in-house collection of heme proteins that displayed activities in other new-to-nature reactions, including various heme-dependent globins and cytochromes *c* (Table 1). Among the hemoproteins examined, *Rhodothermus marinus* nitric oxide dioxygenase (*Rma*NOD) Y32G afforded product **3a** in 39% yield albeit in a racemic manner (Table 1, entry 1). In contrast, a variant of a thermophilic protoglobin from *Aeropyrum pernix, Ape*Pgb W59A Y60G, provided **3a** in lower yield but with promising initial enantioselectivity (24% yield and 77:23 e.r., entry 2). We therefore further evaluated a series of protoglobin variants. Among these, the Y60D F62L V63Q variant of *Thermus amyloliquefaciens* protoglobin (*Tam*Pgb), afforded **3a** in 17% yield and 51:49 e.r. (entry 3). The Y59D F61L V62Q variant of *Thermoanaerobacter* protoglobin (*Tar*Pgb) provided **3a** in 19% yield and 56:44 e.r. (entry 4). The Y57D W59L V60Q variant of *Paracoccus denitrificans* protoglobin (*Par*Pgb) furnished **3a** in 21% yield and 52:48 e.r. (entry 5). Additionally, *Aquifex aeolicus* protoglobin (*Aau*Pgb) Y59D F61L V62Q, and *Pseudomonas fluorescens* protoglobin (*Pfe*Pgb) Y58D W60L V61Q allowed **3a** to be prepared in 24% yield (56:44 e.r., entry 6) and 21% yield (55:45 e.r., entry 7), respectively. Interestingly, our previously engineered *Rhodothermus marinus* thermophilic cytochrome *c* variants for radical C–C of α-diazocarbonyl compounds displayed very low activity and enantioselectivity in the current transformation. For example, *Rma* cyt *c*^RLRDGDE^ (*Rma* cyt *c* V75R M100D M103T T103V M76L T101I M99R T101G V103E)^14^ furnished **3a** in only 3% yield and 47:53 e.r. (entry 8). *Rma* cyt *c*^RIKCGPF^ (*Rma* cyt *c* V75R M100D M103T M76I M99K D100C T101G D102P T103F)^14^ favored the opposite enantiomer, albeit also with low yield and enantiocontrol (5% yield and 52:48 e.r., entry 9). When the yield of the desired product **3a** was low, the homocoupling product derived from **1a** was observed as the major side product. In light of its promising initial activity and enantioselectivity, *Ape*Pgb W59A Y60G was selected as the template for further biocatalyst development for this photobiocatalytic C–C coupling.

**Table 1.** Discovery of photometallobiocatalytic enantioselective C–C coupling of secondary alkyltrifluoroborates and pyridotriazoles^*a*^.

| entry | metalloenzyme | yield (%) | e.r. |
| --- | --- | --- | --- |
| 1 | *RmaNOD* Y32G | 39 | 50:50 |
| 2 | *ApePgb* W59A Y60G | 28 | 77:23 |
| 3 | *TamPgb* Y60D F62L V63Q | 17 | 51:49 |
| 4 | *TarPgb* Y59D F61L V62Q | 19 | 56:44 |
| 5 | *ParPgb* Y57D W59L V60Q | 21 | 52:48 |
| 6 | *AauPgb* Y59D F61L V62Q | 24 | 56:44 |
| 7 | *PfePgb* Y58D W60L V61Q | 21 | 55:45 |
| 8 | *Rma cyt c*<sup>RLRDGDE</sup> | 3 | 47:53 |
| 9 | *Rma cyt c*<sup>RIKCGPF</sup> | 5 | 52:48 |

<sup>a</sup>Reaction conditions: **1a** (4.0 mM, 1 equiv), **2a** (8.0 mM, 2 equiv), 1 mol% metalloenzyme (40 $\mu\text{M}$ ), 10 mol% Eosin Y (400 $\mu\text{M}$ ), $h\nu$ (440 nm), 200 mM KPi buffer (pH = 7.4), rt, 12 h. Protein structures were generated based on their AlphaFold3 model.<sup>20</sup> The yield of **3a** was calculated based on benzyltrifluoroborate salt **1a** as the limiting reagent.

To further enhance the efficiency of this photobiocatalytic C–C coupling, we next evaluated a series of photocatalysts (Table 2). Transition-metal-based photocatalysts^1b^ such as Ru(bpy)_3_Cl_2_, *fac*-Ir(ppy)_3_, and [Ir(ppy)_2_(dtbbpy)]PF_6_ afforded **3a** in low yield (entry 1-3). When [Ir(dF(CF_3_)ppy)_2_(dtbbpy)]PF_6_ was used, no product was detected (entry 4). We next evaluated common organic photocatalysts^1c^ for this asymmetric radical coupling. 4CzIPN afforded **3a** in only 4% yield and 82:18 e.r. (entry 5), whereas acridinium salt was found to be ineffective for this transformation (entry 6). Rhodamine B and bromocresol were also less effective, providing **3a** in 5% yield with 73:27 e.r. (entry 7) and 4% yield with 76:24 e.r. (entry 8), respectively. Organic dyes from the fluorescein family generally exhibited enhanced efficiency. While the parent fluorescein photocatalyst furnished **3a** in a modest 13% yield with 73:27 e.r. (entry 9), halogenated fluorescein derivatives led to improved reaction efficiency. For example, rose bengal and eosin Y afforded **3a** in 20% yield (73:27 e.r.) and 28% yield (77:23 e.r.), respectively (entries 10 and 11).

**Table 2.**
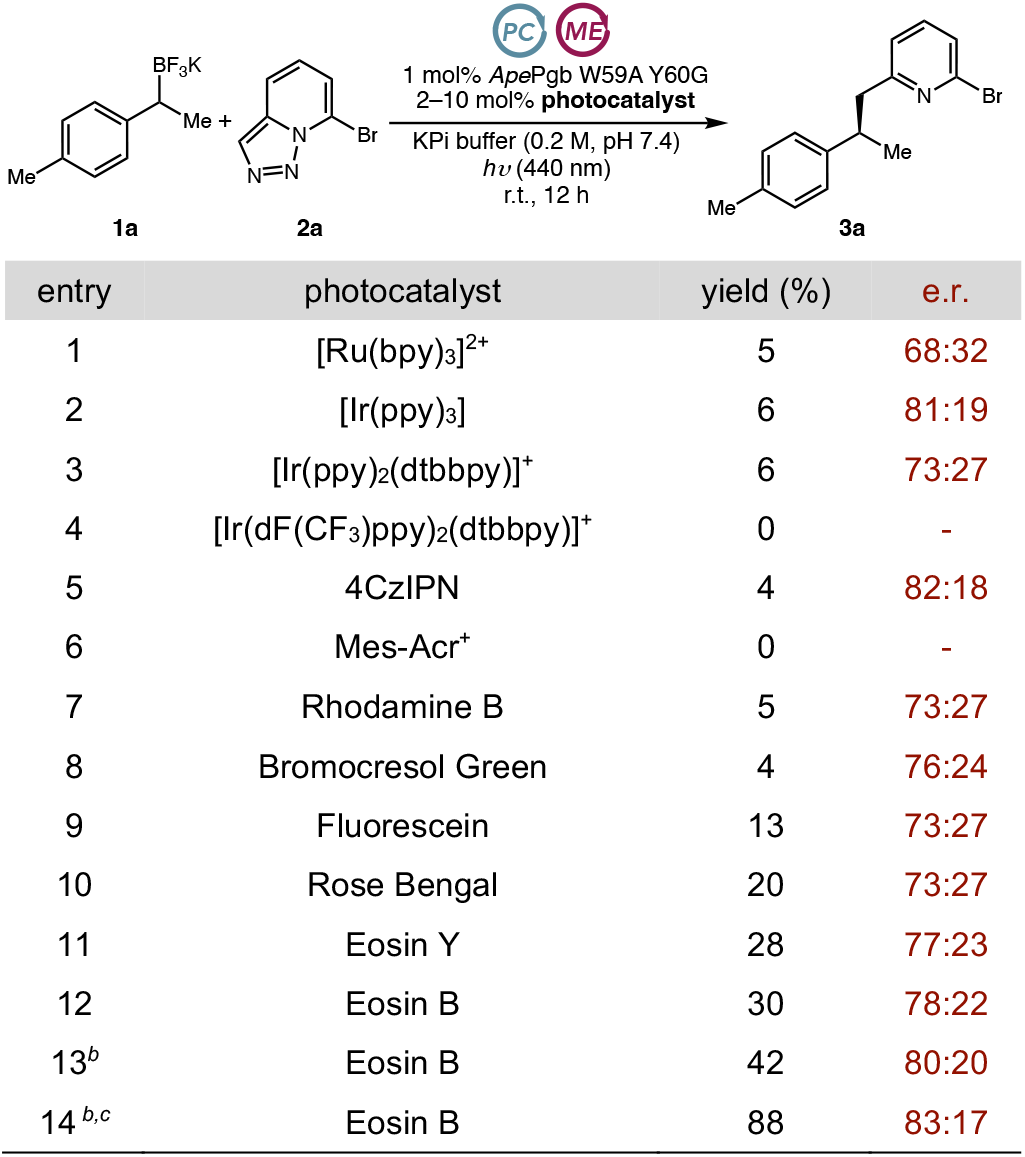
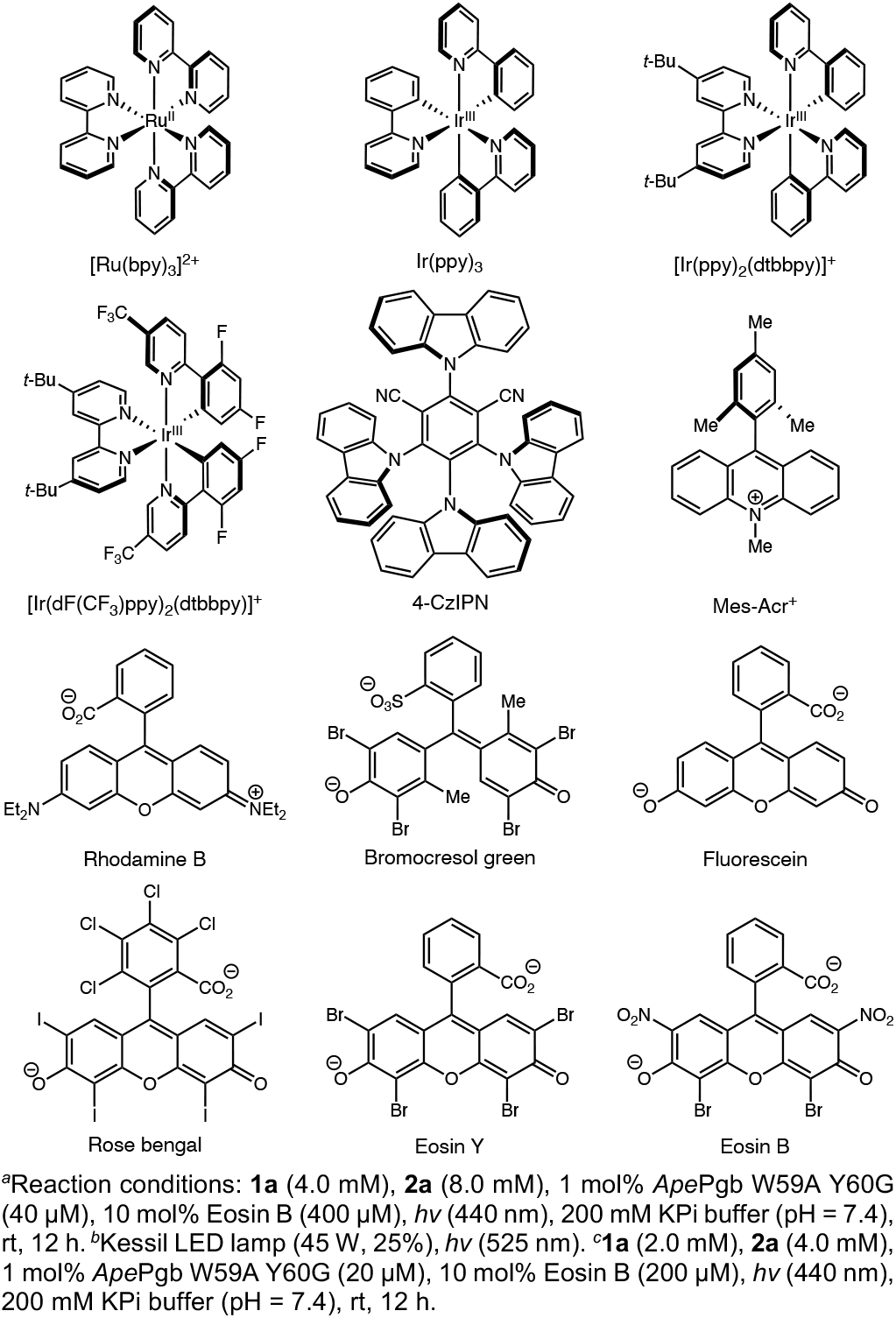
Evaluation of photocatalysts on photometallobiocatalytic enantioselective C–C coupling of secondary alkyltrifluoroborates and pyridotriazoles^*a*^.

| entry | photocatalyst | yield (%) | e.r. |
| --- | --- | --- | --- |
| 1 | [Ru(bpy)<sub>3</sub>]<sup>2+</sup> | 5 | 68:32 |
| 2 | [Ir(ppy)<sub>3</sub>] | 6 | 81:19 |
| 3 | [Ir(ppy)<sub>2</sub>(dtbbpy)]<sup>+</sup> | 6 | 73:27 |
| 4 | [Ir(dF(CF<sub>3</sub>)ppy)<sub>2</sub>(dtbbpy)]<sup>+</sup> | 0 | - |
| 5 | 4CzIPN | 4 | 82:18 |
| 6 | Mes-Acr<sup>+</sup> | 0 | - |
| 7 | Rhodamine B | 5 | 73:27 |
| 8 | Bromocresol Green | 4 | 76:24 |
| 9 | Fluorescein | 13 | 73:27 |
| 10 | Rose Bengal | 20 | 73:27 |
| 11 | Eosin Y | 28 | 77:23 |
| 12 | Eosin B | 30 | 78:22 |
| 13<sup>b</sup> | Eosin B | 42 | 80:20 |
| 14<sup>b,c</sup> | Eosin B | 88 | 83:17 |

<sup>a</sup>Reaction conditions: **1a** (4.0 mM), **2a** (8.0 mM), 1 mol% ApePgb W59A Y60G (40 μM), 10 mol% Eosin B (400 μM), hv (440 nm), 200 mM KPi buffer (pH = 7.4), rt, 12 h. <sup>b</sup>Kessil LED lamp (45 W, 25%), hv (525 nm). <sup>c</sup>**1a** (2.0 mM), **2a** (4.0 mM), 1 mol% ApePgb W59A Y60G (20 μM), 10 mol% Eosin B (200 μM), hv (440 nm), 200 mM KPi buffer (pH = 7.4), rt, 12 h.

Among these organic dyes, eosin B exhibited the highest efficiency, providing **3a** in 30% yield with 78:22 e.r. (entry 12). Given that eosin B displays a maximum absorption wavelength of 535 nm,^14^ we next evaluated the effects of irradiation wavelengths on this photobiocatalytic process. Replacing the blue LED lamp (440 nm) with a green LED lamp (525 nm) further improved both the yield and enantioselectivity of **3a** (42% yield, 80:20 e.r., entry 13). Finally, we examined the effects of light intensity by varying the distance between the LED lamp and the reaction vessel (Table S1). By reducing the light intensity while using eosin B as the photosensitizer, the yield of **3a** could be further improved to 88% with a slightly enhanced e.r. of 83:17 (entry 14).

Using *Ape*Pgb W59A Y60G as the template, we next carried out directed evolution through iterative site-saturation mutagenesis (SSM) and screening to further enhance the enantioselectivity of the protoglobin catalyst in this photobiocatalytic C–C coupling (Figure 1). Active-site residues proximal to the heme cofactor were targeted by iterative saturation mutagenesis and screening. SSM libraries were constructed using the 22c-trick method reported by Reetz.^21^ For each SSM library, 88 clones were selected and evaluated using our in-house high-throughput photobiocatalysis platform consisting of a 96-position parallel photoreactor and a 96-position LED array (530 nm, see the SI for details). Promising variants identified from high-throughput screening with cell-free lysates were subsequently validated in biocatalytic reactions using 1 mol% purified protein catalysts.

**Figure 1.**
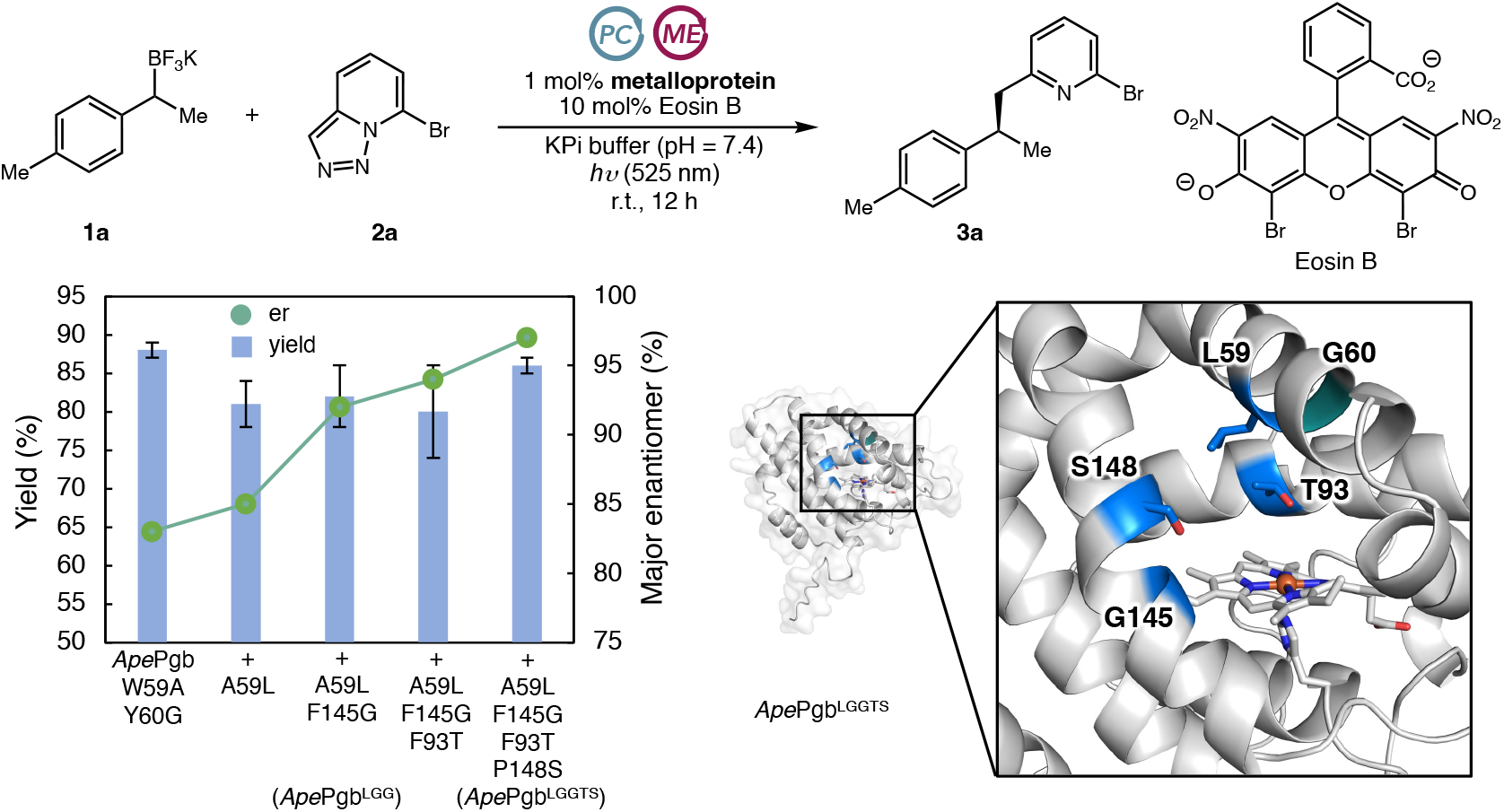
Directed evolution of *Ape*Pgb W59A Y60G for photometallobiocatalytic enantioselective C–C coupling of pyridotriazoles and secondary alkyltrifluoroborate salts. Beneficial mutations discovered from directed evolution are colored in marine. Y60G is colored in deep teal.

Starting from *Ape*Pgb W59A Y60G, SSM targeting A59 afforded a new variant, *Ape*Pgb W59A Y60G A59L (*Ape*Pgb W59L Y60G), which delivered **3a** in (81 ± 3)% yield and an improved enantioselectivity (85:15 e.r.). The second round of directed evolution focused on active-site residues L69, W96, and F145, leading to *Ape*Pgb W59A Y60G A59L F145G, which produced **3a** in (82 ± 4)% yield and 92:8 e.r.. Subsequent SSM targeting residues W62, R90, F93, and I149 afforded *Ape*Pgb W59A Y60G A59L F145G F93T, improving the enantioselectivity of **3a** from 92:8 to 94:6 e.r. A final round of SSM and screening targeting F73, L86, and P148 resulted in the final variant *Ape*Pgb W59L Y60G F145G F93T P148S (*Ape*Pgb^LGGTS^), affording **3a** in (86 ± 1)% yield and 97:3 e.r..

With the evolved *Ape*Pgb^LGGTS^ variant in hand, we next evaluated the substrate scope of this photometallobiocatalytic cross-coupling (Table 3). Organotrifluoroborate salts bearing an *ortho*-(**3b**) and a *meta*-(**3c**) substituent on the aromatic ring were tolerated by *Ape*Pgb^LGGTS^, although slightly lower yields and enantioselectivities were observed. In addition, organotrifluoroborate substrates bearing an electron-donating methoxy (**3d**), an electronically neutral and sensitive methylthio (**3e**), and an electron-withdrawing trifluoromethoxy (**3f**) aryl substituent were readily accommodated under these photobiocatalytic conditions, furnishing enantioenriched pyridine-substituted aliphatic products in good to excellent yields and enantioselectivities. Substrates possessing a halogen substituent, including a fluoro-(**3g**), a chloro-(**3h**), a bromo-(**3i**) and an iodo-(**3j**) group, also underwent the photobiocatalytic C– C coupling to furnish the corresponding products with excellent enantioselectivities. Notably, reactive aryl bromides and iodides were tolerated, demonstrating the mildness of the current reaction conditions. Importantly, preparative scale photobiocatalytic synthesis furnished 124 mg product **3h** in 86% isolated yield without erosion of enantioselectivity relative to analytical scale biotransformations, demonstrating the synthetic value of this new biocatalytic coupling. The trifluoroborate substrate bearing an electron-withdrawing trifluoromethyl substituent at the *meta*-position of the arene (**3k**) could also be successfully converted to the C–C coupling product. Furthermore, a heterocyclic organoboron substrate with a pyridine moiety (**3l**) was transformed with excellent efficiency and enantioselectivity. The organoboron substrate bearing an extended α-ethyl substituent was readily converted, affording the corresponding coupling product (**3m**) with good yield and enantioselectivities. When a more sterically demanding α-propyl organotrifluoroborate substrate was applied, the desired product (**3n**) formed with slightly reduced yield and enantioselectivity. Cyclic secondary alkylboron substrates such as an indanylboron (**3o**) could also be effectively converted with excellent yield and modest enantioselectivity. When six-membered cyclic secondary alkyl radical precursors were used, reduced enantioselectivity was observed with *Ape*Pgb^LGGTS^ despite the excellent yields. We therefore re-examined variants from the *Ape*Pgb^LGGTS^ evolutionary lineage and found that *Ape*Pgb W59L Y60G F145G (*Ape*Pgb^LGG^) served as a highly enantioselective biocatalyst, allowing 1-tetrahydronaphthyl (**3p** and **3q**) as well as chromanyl substrates (**3r**) to be transformed into the corresponding products with excellent yields and enantioselectivities.

**Table 3.**
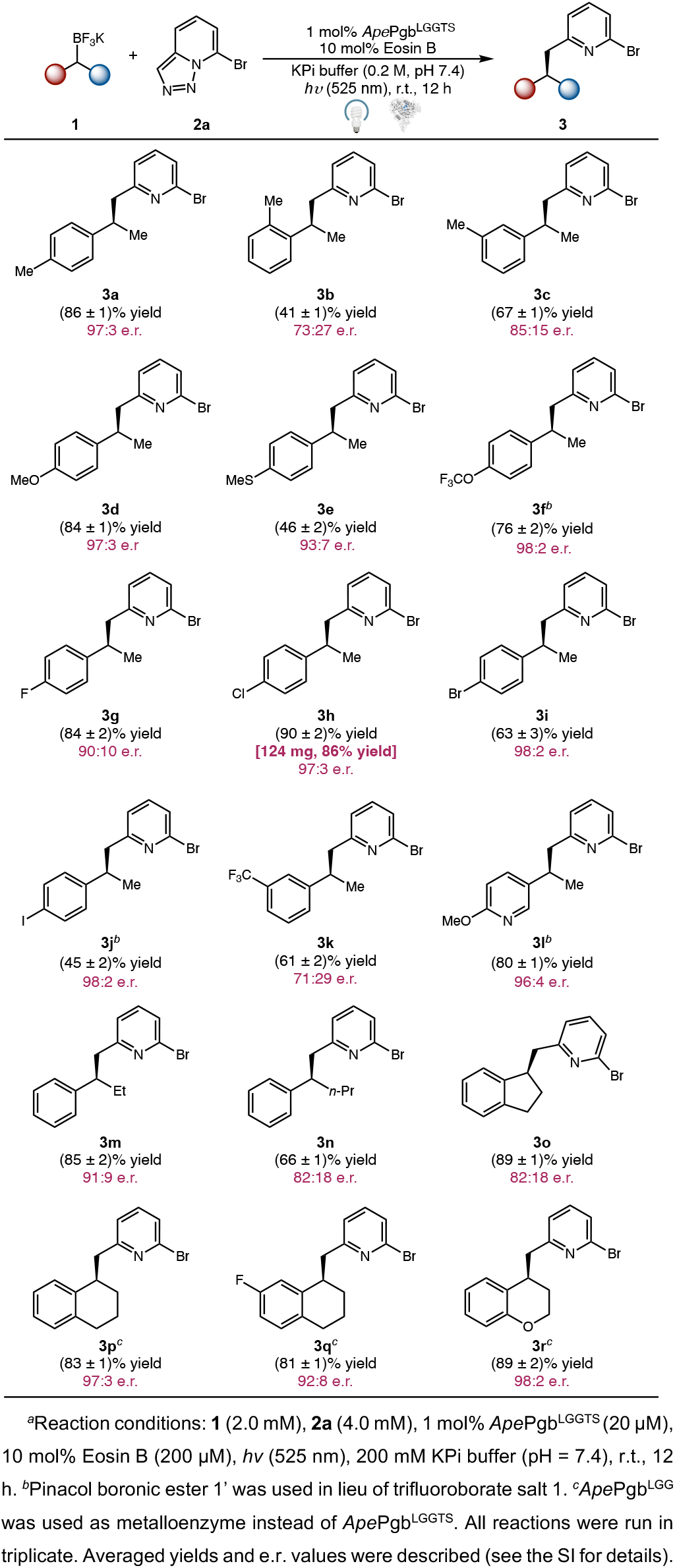
Photometallobiocatalytic enantioselective radical C–C coupling: substrate scope of secondary alkyltrifluoroborate salts^*a*^.

Next, we examined the substrate scope of pyridotriazoles and related nitrogen heterocycles (Table 4). Using *Ape*Pgb^LGGTS^, similar to the 7-bromo substrate (**3a**), 7-chloro-[1,2,3]triazolo[1,5-*a*]pyridine (**3s**) was also successfully converted into the corresponding C–C coupling products with excellent yield and enantioselectivity. 7-iodo-[1,2,3]triazolo[1,5-*a*]pyridine (**3t**) was also a viable substrate, although a reduced yield was observed. Pyridotriazoles possessing a methoxy group (**3u** and **3w**) were also successfully transformed with excellent enantiocontrol. To ascertain the absolute stereochemistry, cross-coupling product **3w** was prepared on a 22 mg scale in 50% isolated yield, and its absolute stereochemistry was assigned by comparison of its specific rotation with literature values.^22^ Finally, a pyrimidine-based heterocycle was also compatible with the current photobiocatalytic coupling protocol, delivering the corresponding coupling product (**3v**) in 84% yield and 83:17 e.r..

**Table 4.**
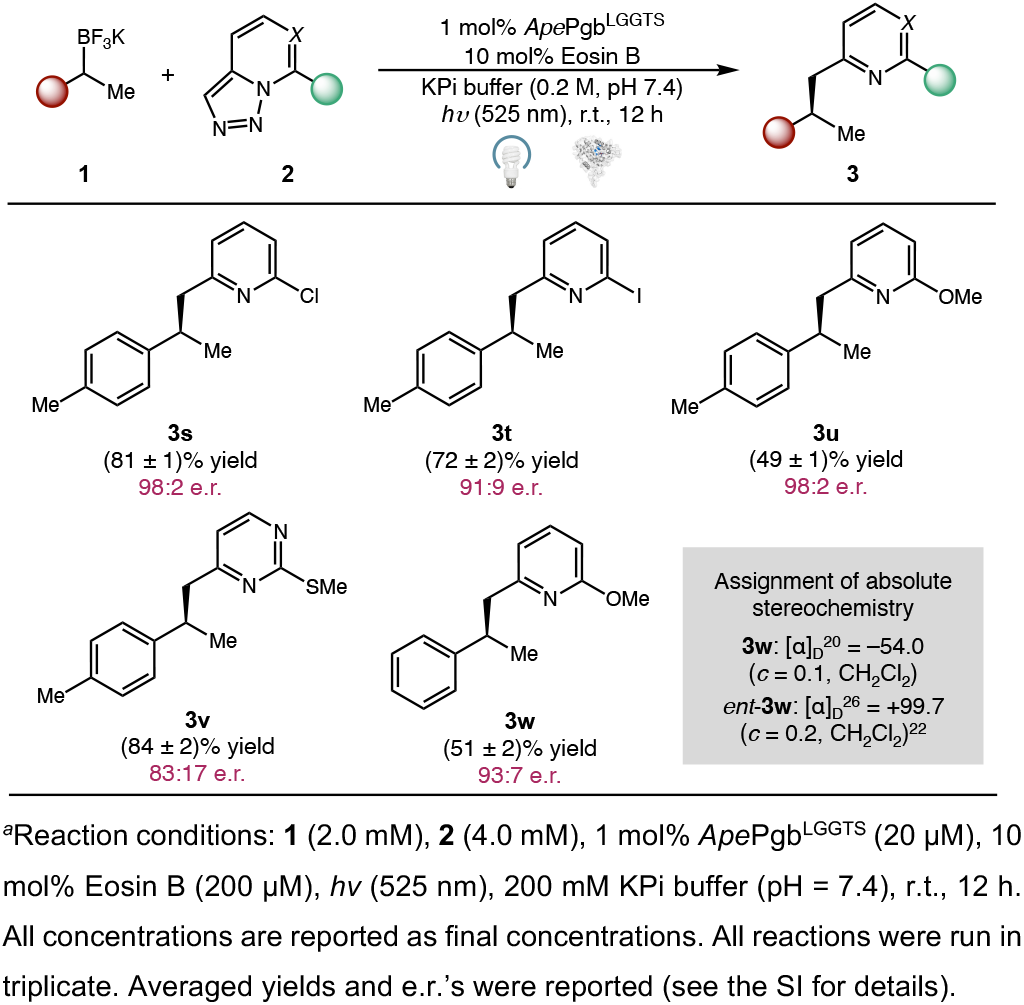
Pyridotriazoles scope of photometallobiocatalytic enantioselective radical C–C coupling^*a*^.

To gain further mechanistic insights into the present photometallobiocatalytic cross-coupling, we conducted a series of experiments (Scheme 2). First, because benzyltrifluoroborate salts are known to undergo protodeborylation under aqueous conditions^23^ to afford the corresponding hydrocarbon 1-ethyl-4-methylalkylbenzene (**4a**), we investigated whether the alkylbenzene **4a** could serve as the precursor to the C–C coupling product **3a** under our conditions (Scheme 2(A)). When compound **4a** was employed in place of benzyltrifluoroborate **1a** under the standard reaction conditions, either in the presence or absence of the photocatalyst eosin B, no coupling product **3a** was detected. This result indicated that the observed C–C bond formation does not arise from a classical Fe-carbenoid-mediated C–H insertion^13c^ pathway involving the protodeborylated hydrocarbon. In addition, under the standard reaction conditions, the heme cofactor itself showed no catalytic activity (Scheme 2(B)), demonstrating the critical role of the protein scaffold in enabling the newly developed photobiocatalytic C–C coupling. Furthermore, radical trapping experiments were performed using 5.0 equiv 2,2,6,6-tetramethylpiperidin-1-oxyl (TEMPO) (Scheme 2(C)). In the presence of TEMPO, the desired photobiocatalytic coupling product **3a** was not observed while the corresponding TEMPO trapping adduct **5a** was obtained in 48% yield, supporting the involvement of the corresponding secondary radical intermediate generated under the photobiocatalytic reaction conditions.

**Scheme 2.**
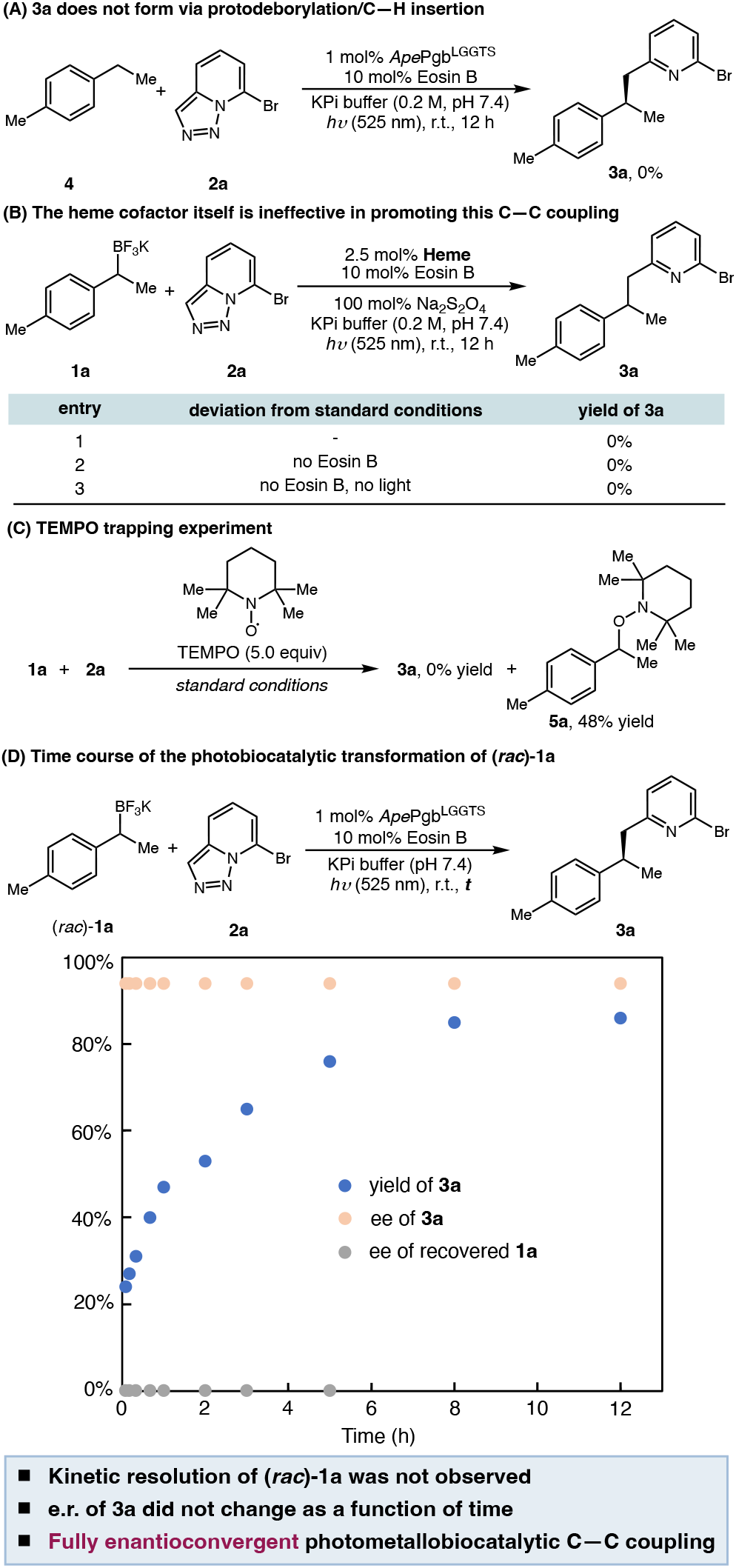
Photometallobiocatalytic asymmetric cross-coupling of benzyltrifluoroborate salts and pyridotriazoles^*a*^ PC = photoredox catalyst, ME = metalloenzyme.

To interrogate the stereochemical course of the photobiocatalytic C–C coupling, time-course analysis was performed for the conversion of (*rac*)-**1a** into enantioenriched **3a** (Scheme 2(D)). The photobiocatalytic reaction was quenched at defined time intervals, and aliquots of the reaction mixture were withdrawn to determine the conversion of **1a**, the yield of **3a**, and the enantiomeric excess of **1a** and **3a**. It was found that the e.r. of recovered **1a** remained 50:50 throughout the transformation, indicating that no kinetic resolution of **1a** occurred throughout the transformation. This finding is consistent with radical generation outside the enzyme active site. Additionally, the e.r. of the product **3a** remained constant over time, thereby establishing the enantioconvergent nature of this photobiocatalytic cross-coupling from racemic organoboron substrates. Together, these results are also consistent with our proposed radical-mediated mechanism allowing enantioconvergent transformations of secondary alkylboron substrates.

To probe the interactions between the protoglobin biocatalyst and the photocatalyst eosin B, we carried out fluorescence anisotropy titration using a plate reader-based high-throughput assay (Figure 2).^24^ The fluorescence polarization titration data were fit using a single-site binding model to determine the equilibrium dissociation constant (*K*_d_) between eosin B and *Ape*Pgb^LGGTS^. These measurements revealed that eosin B forms a complex with *Ape*Pgb^LGGTS^, with a *K*_d_ of (5.4 ± 0.8) μM, indicating spontaneous binding of eosin B to the engineered metalloenzyme. Under the photobiocatalytic reaction conditions, this *K*_d_ indicates that 97% *Ape*Pgb^LGGTS^ is bound to eosin B. We postulate that the spontaneous association of the biocatalyst and the photocatalyst may play a role in accounting for the excellent activity of eosin B under the optimized photobiocatalytic conditions.

**Figure 2.**
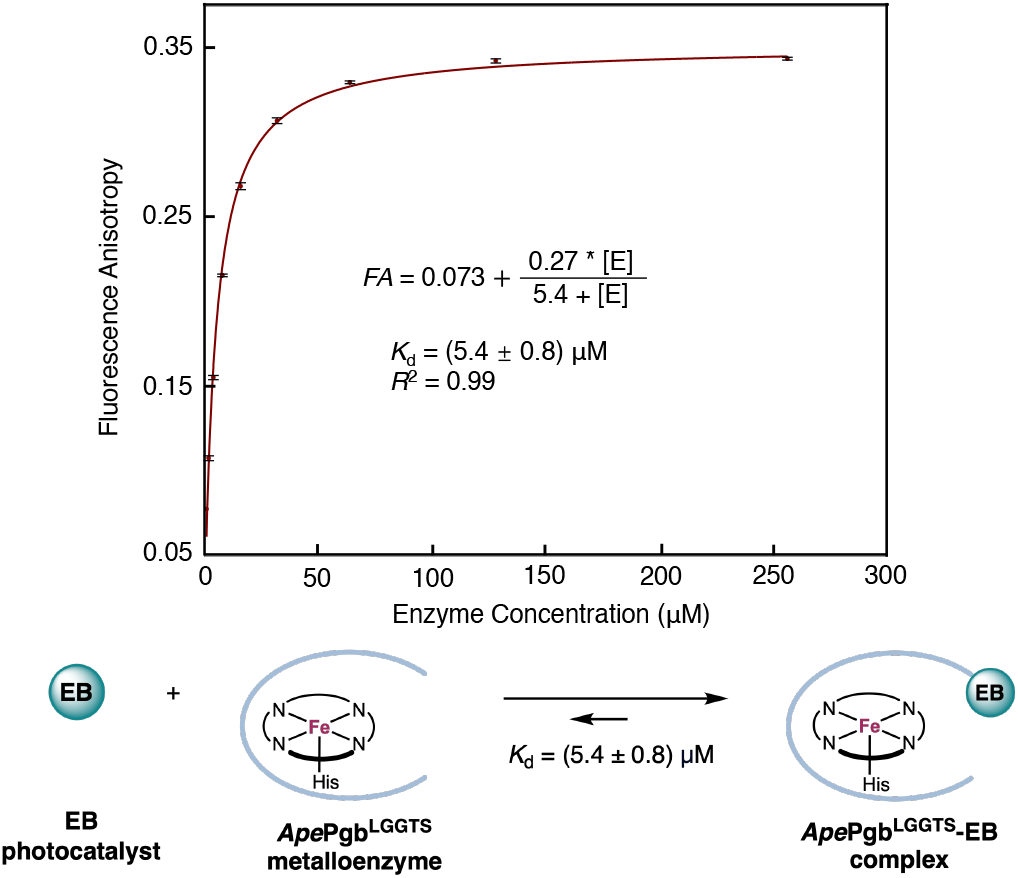
Fluorescence anisotropy titration measurements with *Ape*Pgb^LGGTS^ and eosin B.

### Computational Studies

To elucidate the effects of beneficial mutations introduced through directed evolution on the enantioselectivity in the radical-mediated C–C bond formation step, classical molecular dynamics (MD) simulations were performed to model the near-attack conformations (NACs) for the S_H_2 reaction between the secondary benzylic radical and the Fe–porphyrin alkyl intermediate in *Ape*Pgb^LGGTS^ (Scheme 1(B); see Supporting Information for Computational Details). In these NAC simulations, the distance between the benzylic carbon of the radical and the Cα of the Fe-alkyl species was restrained to a range of 2.7–2.9 Å, based on our DFT-optimized transition state geometry (Figure S15). MD simulations show that the beneficial mutations F145G and P148S induce a pronounced conformational rearrangement of the α-helix spanning residues 139–160, positioning residue I149 closer to the Fe–alkyl intermediate (Figure S16). In the disfavored (*Re*)-face attack pathway, I149 forms steric clash with the methyl group of the benzylic radical with an average H···H distance of 2.50 ± 0.28 Å, which destabilizes the NAC for S_H_2 substitution at the (*Re*)-face of the prochiral radical (Figures 3(B) and S18). In addition, the W59L mutation introduces close contacts between L59 and the methyl group of the benzylic radical (2.51 ± 0.27 Å), further destabilizing the (*Re*)-face substitution NAC. These unfavorable steric repulsions are alleviated in the preferred (*Si*)-face C–C bond formation, as evidenced by the longer methyl–W59L and methyl–I149 distances of 2.60 ± 0.34 Åand 3.04 ± 0.70 Å, respectively (Figures 3(A) and S17). Additionally, the phenyl group of the benzylic radical is positioned proximal to residue G60, suggesting that the reduced steric bulk at this position provides additional space to accommodate the radical and, thus, facilitates the C–C bond formation. Taken together, our classical MD simulations reveal how beneficial mutations reshape the enzyme active site to selectively destabilize the (*Re*)-face substitution NAC, providing the structural basis for the observed enantioselectivity in the radical C–C bond formation.

**Figure 3.**
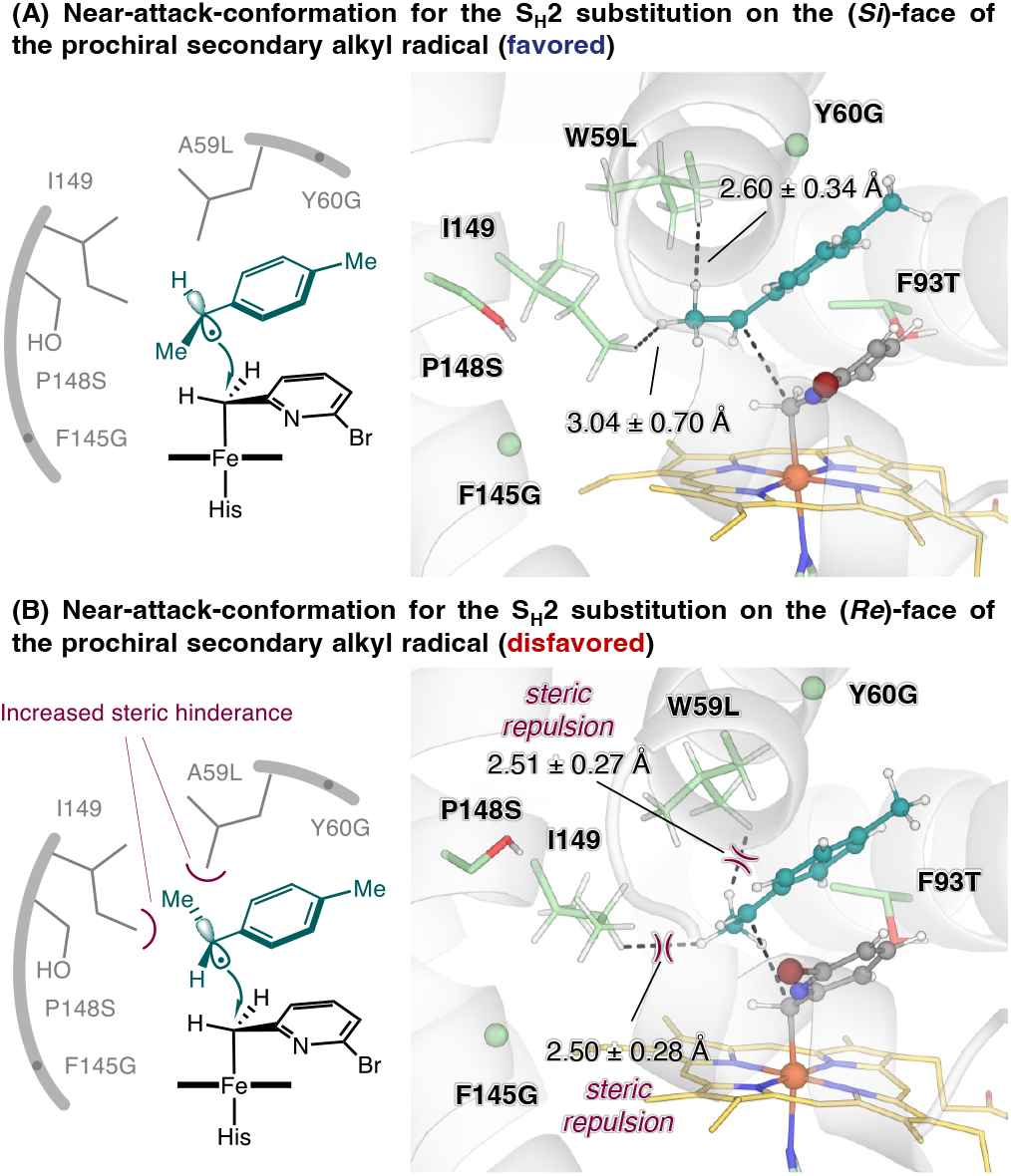
Origin of enzymatic enantiocontrol in the *Ape*Pgb^LGGTS^-catalyzed radical cross-coupling of a prochiral secondary alkyl radical revealed from classical MD simulations.

## Conclusion

In summary, we developed a cooperative photometallobiocatalytic strategy for the enantioselective intermolecular C–C cross-coupling of pyridotriazoles and secondary alkyltrifluoroborate salts through synergistic catalysis between an organic photocatalyst and an engineered protoglobin. Directed evolution of thermostable *Aeropyrum pernix* protoglobin furnished highly efficient biocatalysts capable of enabling stereocontrolled radical cross-coupling reactions involving enzymatic iron carbenoid intermediates. The resulting photobiocatalytic platform accommodates a broad range of organotrifluoroborate salts and pyridotriazoles, providing access to structurally diverse *N*-heterocyclic products with excellent yields and enantioselectivities. Mechanistic studies support the intermediacy of secondary alkyl radicals and suggest that spontaneous association between eosin B and the engineered metalloenzyme may facilitate productive radical delivery during catalysis. Collectively, these results demonstrated an emerging outer-sphere mechanism for stereoselective formal metal carbenoid–radical cross-coupling. Beyond expanding the reactivity of metal carbenoid chemistry, this work highlighted the potential of cooperative photometallobiocatalysis to enable new classes of asymmetric radical transformations that are challenging to achieve using conventional small-molecule catalysis or biocatalysis alone.

## Acknowledgements

We are grateful to the National Institutes of Health (R35GM147387) for financial support. Computational study is supported by the National Science Foundation (CHE-2400087 to P.L.). Y.Y. is an Alfred P. Sloan Research Fellow (FG-2024-22244), a Camille Dreyfus Teacher-Scholar Awardee (TC-25-084), a David & Lucile Packard Fellow (2023-76169) and a Howard Hughes Medical Institute Freeman Hrabowski Scholar. Computations were carried out at the University of Pittsburgh Center for Research Computing and Data and the Advanced Cyberinfrastructure Coordination Ecosystem: Services & Support (ACCESS) Program through allocations CHE-140139 and CHE-260005, which is supported by NSF awards OAC-2117681, OAC-1928147, OAC-1928224, and CHE-260005.

